# Talin Folding as the Tuning Fork of Cellular Mechanotransduction

**DOI:** 10.1101/2020.01.10.901991

**Authors:** Rafael Tapia-Rojo, Alvaro Alonso-Caballero, Julio M. Fernandez

## Abstract

Cells continually sample their mechanical environment using exquisite force sensors such as talin, whose folding status triggers mechanotransduction pathways by recruiting binding partners. Mechanical signals in biology change quickly over time and are often embedded in noise; however, the mechanics of force-sensing proteins have only been tested using simple force protocols, such as constant or ramped forces. Here, using our magnetic tape head tweezers design, we measure the folding dynamics of single talin proteins in response to external mechanical noise and cyclic force perturbations. Our experiments demonstrate that talin filters out external mechanical noise but detects periodic force signals over a finely-tuned frequency range. Hence, talin operates as a mechanical bandpass filter, able to read and interpret frequency-dependent mechanical information through its folding dynamics. We describe our observations in the context of stochastic resonance, which we propose as a mechanism by which mechanosensing proteins could respond accurately to force signals in the naturally noisy biological environment.

## INTRODUCTION

The precise interpretation of complex mechanical signals underpins a vast range of biological processes [1–3]. These force signals are accurately detected by cellular force sensors and transduced into electrical or biochemical signals to trigger a physiological response—mechanotransduction [4, 5]. For instance, the human auditory system converts complex vibration patterns into electrophysiological signals with astonishing speed and accuracy, detecting frequencies from 20 to 20,000 Hz with a sensitivity finer than the music interval and a dynamic range greater than 120 dB [6, 7]. Akin to sound waves, mechanical signals in biology oscillate at frequencies that span over several timescales and, yet, are effectively decomposed and interpreted by cells. Furthermore, in addition to the intrinsic thermal noise that drives Brownian motion, athermal noise is a protagonist at the molecular and cellular level [8]. Every mechanical communication is thus challenged by a background of non-informative fluctuations that, still, do not interfere with a robust cellular response. However, how molecular force sensors interpret complex mechanical signals in a finely-tuned way remains mostly unknown.

Cells continuously receive and transmit mechanical signals between neighbor cells and the extracellular matrix, which guides diverse processes such as embryonic development [9], cell traction [10], or tumor proliferation [11]. These force stimuli are transduced into biochemical signals, generally by inducing conformational changes in force-bearing proteins that modulate the affinity of binding partners or induce posttranslational modifications [4, 12, 13]. For instance, cells migrate following the stiffness gradient of their substrate, which is sampled by periodic forces applied by focal adhesions—durotaxis [14, 15]. Talin is a mechanosensing hub protein in focal adhesions, which crosslinks transmembrane integrins with the active F-actin filaments and recruits several binding proteins to control the function and fate of this organelle [16–18]. For example, vinculin binds to cryptic helices in mechanically unfolded talin domains, subsequently recruiting actin filaments that reinforce the cellular junction [13, 19–21]. Hence, talin transduces mechanical forces through its folding dynamics.

The mechanics of talin have been studied using force spectroscopy techniques, which typically apply ramped or constant forces to monitor folding or unfolding reactions [22, 23]. These studies have shown that force drives talin response by unfolding each of the 13-rod bundle domains hierarchically; increasing forces result in a gradual unfolding of the talin rod [24, 25]. This response establishes a simple mechanotransduction pathway; the force load on talin determines the number of unfolded talin domains and, hence, how many vinculin molecules can be recruited. However, the response of talin under complex mechanical signals has never been studied, so it remains unknown how talin folding dynamics couple to forces that change in time or if its response is affected when the mechanical perturbation is buried in external noise fluctuations.

Here, we measure the folding dynamics of the talin R3 domain under complex force signals, which include external mechanical noise and sinusoidal perturbations. Using our new magnetic tape head tweezers design (Fig. 1 A), we demonstrate that talin rejects external mechanical noise but exhibits phase-locked folding dynamics when subject to oscillating forces. Talin response is finely tuned, and it only detects signals over a narrow frequency range, which is defined by its mechanical stability; the R3 WT domain responds to signals around 7 Hz, while the more stable R3 IVVI mutant detects slower signals at 1 Hz. We ascribe our results to the physical phenomenon of stochastic resonance, which we propose as a mechanism by which force-sensing proteins could interpret complex mechanical signals through their folding dynamics, by operating as a “biochemical Fourier transform.”

**FIG. 1.**
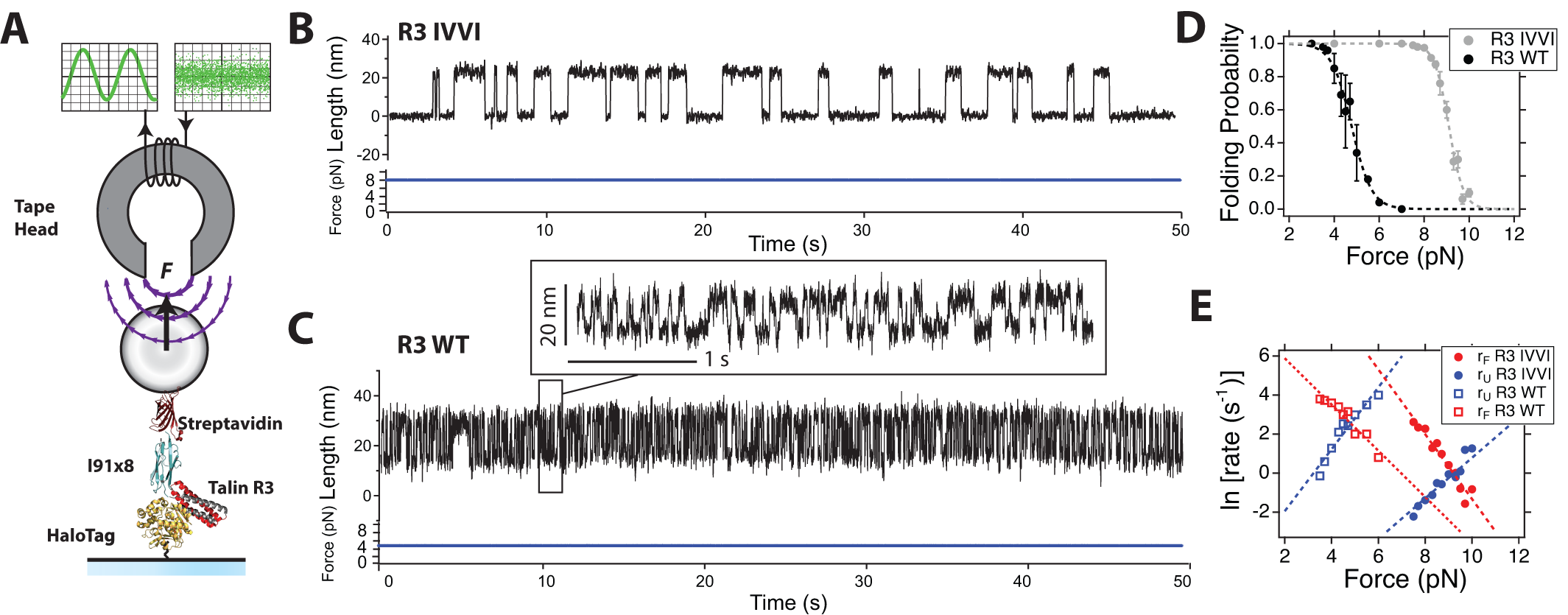
Mechanical characterization of the talin R3 domain: (A) Schematics of our magnetic tweezers assay for applying complex force signals to single proteins. Our construct contains the talin R3 domain followed by eight repeats of the titin I91 domain, flanked by a HaloTag for covalent anchoring to the glass surface and biotin for tethering with streptavidin-coated M-270 magnetic beads. Forces with a bandwidth greater than 10 kHz are generated using a magnetic tape head. (B) Typical single-molecule recording of talin R3 IVVI at 9 pN. (C) Typical recording of talin R3 WT at 5 pN. (Inset) Detail of the trajectory, highlighting the faster dynamics of R3 WT compared to the mutant R3 IVVI. Traces acquired at a frame rate between 1 and 1.5 kHz and smoothed with a Savitzky-Golay 4th-order filter, with a 101-points box (R3 IVVI) and a 31-points box (R3 WT). (D) Folding probability of the R3 IVVI and R3 WT talin domains. The R3 IVVI mutant has a higher mechanical stability, and its folding probability is shifted to higher forces by 4 pN. (E) Folding (red) and unfolding (blue) rates for R3 IVVI (solid) and R3 WT (empty) talin domains. At the coexistence force, the rates of the WT are ∼12 s^−1^, compared to the slower ones of R3 IVVI (∼1.5 s^−1^). Data fitted with the Bell-Evans model (dotted lines), yielding 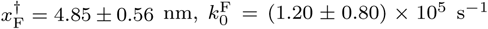 and 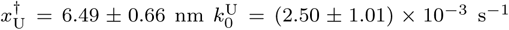 for R3 WT, and 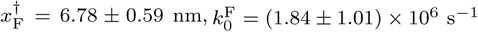 and 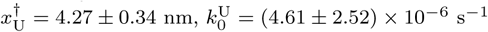 for R3 IVVI.

## RESULTS

### Mechanical Characterization of the Talin R3 Domain

The talin R3 domain is the weakest of all 13-rod domains, thought to be the precursor of talin-based adhesions, given its low unfolding force and the presence of two vinculin-binding sites [16, 21, 24, 25]. This low mechanical stability is due to the four-residue threonine belt on its hydrophobic core; the substitution of these residues in the mutant R3 IVVI increases its stability significantly [16, 21]. Due to these different mechanical properties, we explore both the R3 WT and R3 IVVI domains perturbed by complex mechanical signals, which will allow us to compare and relate their responses with their mechanics. First, in order to establish the benchmarks for the study under complex force signals, we characterize the folding mechanics of both domains at constant force (Fig. 1).

Figures 1B and C show representative recordings measured at the coexistence force (F_1*/*2_; same occupation of the folded and unfolded states), which is 9 pN for R3 IVVI, and 5 pN for R3 WT. Their folding dynamics are characterized by reversible transitions between the folded and unfolded states with an extension change of ∼20 nm, but the kinetics of R3 WT is nearly 10 times faster, compared to those of R3 IVVI (Fig. 1C, inset). From equilibrium trajectories as these, we measure the relative occupation of the folded state (folding probability; Fig. 1D), and the folding (r_F_) and unfolding (r_U_) rates (Fig. 1E). The folding probability spans less than 2 pN, illustrating the exquisite force sensitivity of both domains. Due to this steep force dependency, the folding and unfolding rates have an exponential Bell-like dependency 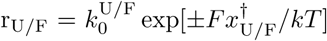 [26]), with similar distances to transition state from the folded and unfolded states. Overall, both domains exhibit two-state like dynamics, occurring at different force ranges— between 4 and 6 pN (R3 WT), and 8 to 10 pN (R3 IVVI)—and kinetic rates on a very different timescale—at F_1*/*2_, r_*F/U*_ ≈12 s^−1^ (R3 WT), and r_*F/U*_ ≈1.5 s^−1^ (R3 IVVI).

### Talin Discerns between External Mechanical Noise and Oscillatory Forces

Motivated by the naturally noisy cellular environment, we first asked if talin maintains a robust response when subject to a randomly fluctuating perturbation instead of a constant force. Applying forces that change rapidly is challenging, as force spectrometers typically change the force by displacing large mechanical probes. To this aim, we take advantage of our new magnetic tape head tweezers design, able to change the force with a bandwidth greater than 10 kHz while maintaining intrinsic force-clamp conditions (Fig. 1A, [27]). We generate external mechanical noise as a broadband uncorrelated force signal, with an average of 9 pN and a peak-to-peak amplitude greater than 7 pN (SI Appendix, Fig. S1 and Supplementary Methods) and apply it to the talin R3 IVVI domain. This signal fluctuates around *F*_1/2_ over a force range that significantly exceeds the folding regime of R3 IVVI. Importantly, this external mechanical noise is different from the intrinsic thermal noise provided by the heat bath, which we do not control in our experiment and that induces the stochastic folding and unfolding transitions we observe at constant force.

Figure 2A (left) shows the dynamics of R3 IVVI perturbed by external mechanical noise. Strikingly, despite the fine force sensitivity of talin and the broad force range over which the external mechanical noise fluctuates, we do not observe any effect on talin dynamics, which are statistically indistinguishable from those measured at the average force of 9 pN. The dwell-time histograms—obtained using a maximum-likelihood procedure (SI Appendix, Fig. S2)—are both exponential distributions with approximately the same time constant—0.59 *±* 0.02 s (noise) versus 0.69 *±* 0.02 s (no noise) (Fig. 2A, right). This apparent insensitivity to external mechanical fluctuations is maintained at any other folding force. We observe no statistically significant differences (within a 95% confidence interval) between the folding probabilities measured in the presence or absence of external mechanical noise, and the coexistence forces (F_1*/*2_) obtained under both conditions overlap within error bars (SI Appendix, Fig. S3). This evidence suggests that talin has a robust mechanical response, filtering out the randomly fluctuating components, and responding only to the average force.

**FIG. 2.**
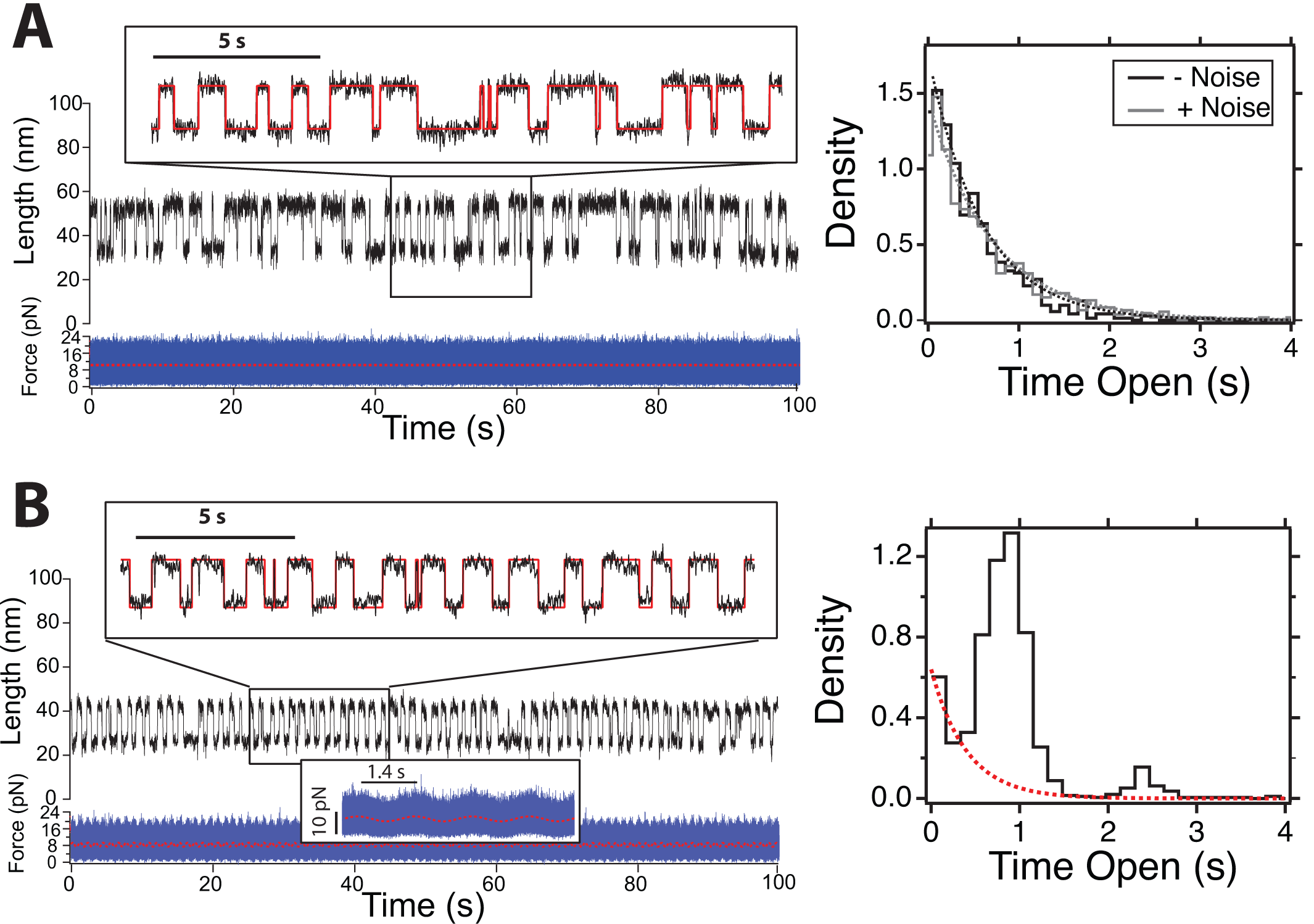
Talin detects oscillatory signals among mechanical noise: (A) (Left) Dynamics of the talin R3 IVVI domain under mechanical white noise with an average of 9 pN and a standard deviation of 2.8 pN. (Inset) Detail of the trajectory with the idealized trace (red). (Right) Comparison of the dwell-time histograms in the unfolded state perturbed by a constant force of 9 pN (grey) and external mechanical noise (black). Both distributions are comparable, as talin remains insensitive to randomly fluctuating forces and responds only to the average perturbation. Histograms built from 1063 (no noise) and 704 transitions (noise). (B) (Left) Dynamics of the talin R3 IVVI domain perturbed by a force signal oscillating at 0.7 Hz, with an amplitude of 1 pN, and submerged in mechanical white noise with a standard deviation of 2.8 pN. (Right) Dwell-time histogram measured under such signal. The folding transitions are synchronized with the oscillating force, while the mechanical noise is ignored. These dynamics shift the shape of the dwell-time distribution, which shows resonance peaks at odd multiples of half the period of the signal. The remnant stochastic behavior is appreciated as an underlying exponential distribution (red dotted line). Histogram built with 2295 transitions. Traces acquired at a frame rate of 1-1.5 kHz, and smoothed with a Savitzky-Golay 4th order algorithm with a 101-points box size.

Given the ability of talin to remain indifferent to the external mechanical noise, we explore its response under a force signal that oscillates at a specific frequency and is submerged in external mechanical noise. We build this signal by adding a sinusoidal force *F* (*t*) = *F*_0_ +*A* sin(2*πf*_Ω_) (with *F*_0_ = 9 pN, *A* = 1 pN and *f*_Ω_ = 0.7 Hz) to external mechanical noise with the same properties employed before. Hence, the amplitude of the correlated component is small compared to the background fluctuations (Fig. 2B, force inset). By contrast, talin dynamics are now dramatically altered, and most transitions occur at regular intervals of ∼0.7 s, while the mechanical noise is ignored (Fig. 2B, left). These entrained folding dynamics shift the shape of the dwell-time histogram from an exponential to a peaked distribution (Fig. 2B, right). The first peak is centered at ∼0.7 s, which coincides with half-period of *f*_Ω_, while the second peak appears at three times that value, ∼2.1 s. Despite the dominant entrained dynamics, we observe an underlying exponential distribution, which reflects a minority of remnant stochastic transitions, occurring at a time constant compatible to a force of 9 pN (Fig. 2B, right, red dotted line). Hence, unlike random force fluctuations, talin detects force oscillations through its folding dynamics, which shift from stochastic—characterized by exponentially-distributed dwell-times—, to a phase-locked response, reflected in peaked dwell-time distributions.

### Mechanical Signal Detection by Talin is Frequency-Dependent

The talin R3 domain detects minimal force changes through its folding dynamics; differences in force of only 0.3 pN shift the folding probability by up to 30% (Fig. 1D). Since our data demonstrates that talin detects oscillatory forces, we seek to test if such a response is also frequency-sensitive. To this aim, we subject R3 IVVI to purely oscillating forces at different frequencies, *f*_Ω_ = 0.1, 1, and 10 Hz, with a mean value of *F*_0_ = 9 pN, and amplitude of *A* = 0.7 pN. Figure 3 shows R3 IVVI folding dynamics under these three oscillatory force signals (left), with the associated dwelltime histograms (right). Under the slowest perturbation (*f*_Ω_ = 0.1 Hz; Fig. 3A), we observe that, while the extension of the protein follows the oscillation, most transitions are stochastic as the force changes much more slowly than the natural folding kinetics—0.1 Hz versus ∼1.5 Hz. The dwell-time histogram is an exponential distribution with a time constant compatible to a constant force of 9 pN (*r*_K_ = 2.16 *±* 0.15 s^−1^), and a marginal contribution of entrained transitions as a small peak at 5 s (red star). At *f*_Ω_ = 1 Hz (Fig. 2B), most folding transitions are synchronized with the perturbation and occur at regularly spaced time intervals. The dwell-time histogram has three peaks of diminishing amplitude and a small underlying exponential distribution. These peaks are centered at odd multiples of half the driving period—0.5, 1.5, and 2.5 seconds—and represent harmonics of talin phase-locked response. Finally, at a driving frequency of 10 Hz (Fig. 3C), the stochastic behavior is recovered with the same dynamics as those measured at 9 pN. The dwell-time histogram is an exponential distribution with a time constant of *r*_*K*_ = 2.13 *±* 0.10 s^−1^, which indicates that talin filters this fast oscillation. Only when calculating the dwell-time distribution with logarithmic binning, we observe a small contribution at 0.05 s corresponding to a minority of entrained transitions (Fig. 3C, inset). Hence, these data indicate a frequency-selective response of talin upon oscillating forces. Interestingly, talin folding dynamics segregate into phase-locked dynamics that respond to the oscillation with a frequency-dependent intensity, and a stochastic behavior that filters the signal and detects only the average of the perturbation.

**FIG. 3.**
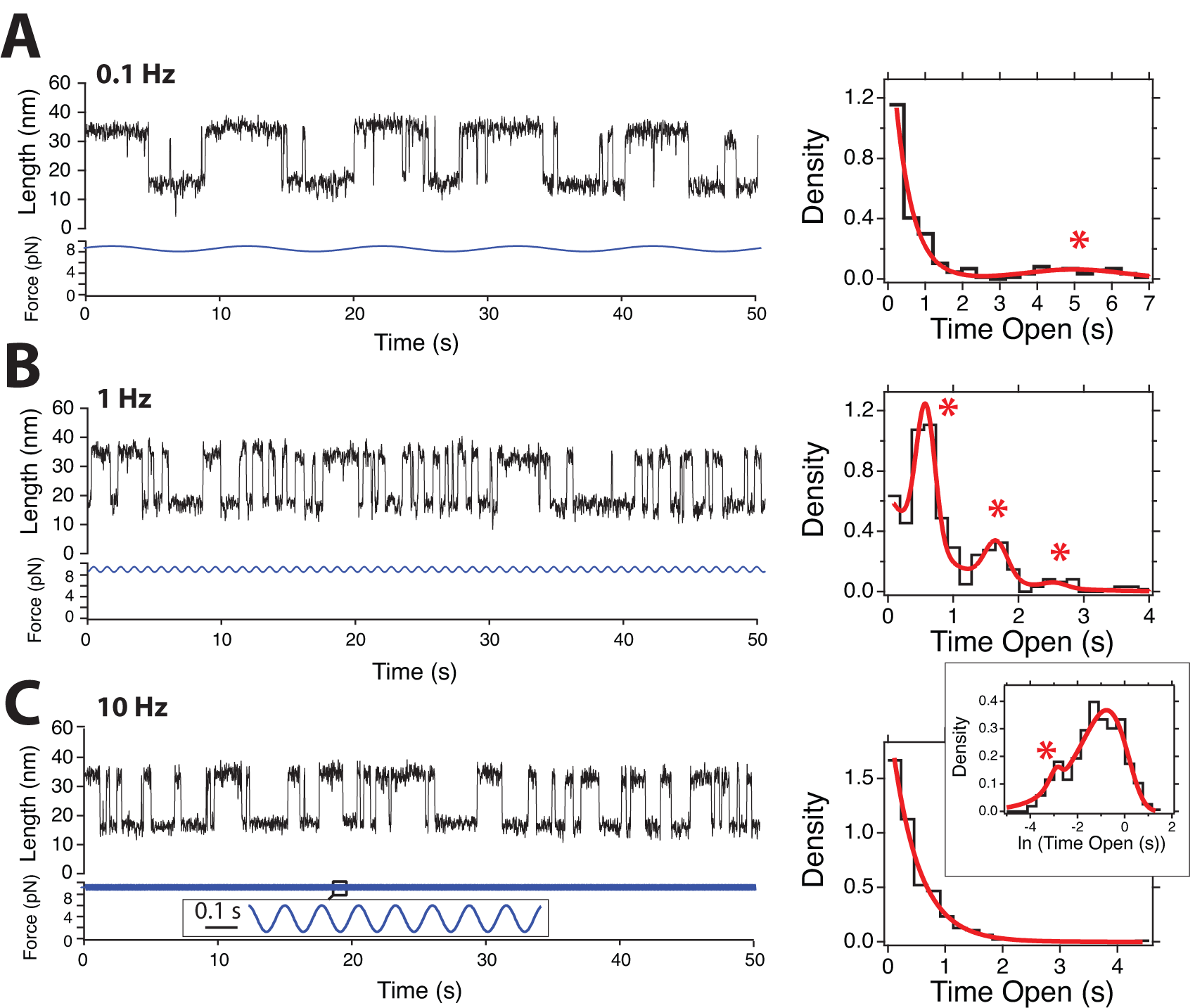
Dynamics of the talin R3 IVVI domain under signals of different frequencies: (A, B, C) (Left) 50 seconds-long fragments of magnetic tweezers recordings showing R3 IVVI under a periodic signal with an amplitude of 0.7 pN, and frequencies of 0.1, 1, and 10 Hz, respectively. (Right) Dwell-time histograms on the unfolded state calculated for each of these signals. The interplay between the stochastic and resonant dynamics is manifested as a combination of exponential and Gaussian distributions, with relative weights that indicate the level of response (red fits). At 0.1 Hz, most transitions are stochastic, and only a small peak at 5 s appears (red star). At 1 Hz, the response is optimal and the majority of the histogram is built by Gaussian peaks, located at odd multiples of half-period of the driving signal (red stars). At a high frequency of 10 Hz, the distribution is exponential, and only a logarithmic binning reveals a small proportion of resonant transitions at 0.05 s (inset, red star). Histograms built with 223 (A), 337 (B) and 363 (C) transitions. Traces acquired at a frame rate between 1 and 1.5 kHz, and smoothed with a Savitzky-Golay 4th-order filter, with a 101-points box.

### Talin Response is Governed by the Mechanical Stability of the Domain

The response of the R3 IVVI domain suggests that the entrained dynamics manifest optimally at frequencies of the order of the natural folding rates, which at 9 pN are ∼1.5 s^−1^ (Figure 1E). Hence, it can be expected that proteins with faster kinetic rates will respond to higher frequencies and vice versa, providing a tuning mechanism based on the mechanical stability of the protein domain. In order to test this hypothesis, we measure the folding dynamics of the R3 WT domain subject to oscillatory forces given that, at the coexistence force *F*_1*/*2_ = 5 pN, its kinetic rates are nearly 10 times faster (∼12 s^−1^; Fig. 1E).

Figure 4 shows single-molecule recordings of R3 WT perturbed with sinusoidal signals of 0.5 (A), 5 (B), and 50 Hz (C), an average of *F*_0_ = 5 pN, and amplitude of *A* = 0.7 pN. These data show an analogous response to that measured for R3 IVVI but shifted to higher frequencies. At 0.5 Hz, the dynamics are exponentially distributed, with a minor entrained contribution appreciable in log-scale (Fig. 4A, inset). At 5 Hz, the transitions are entrained with the force signal, showing a peak at 0.1 s and an underlying exponential distribution with a time constant compatible to that measured at 5 pN (*r*_*K*_ = 18. *±* 0 2.24 s^−1^). At 50 Hz, the dynamics are stochastic with a rate of *r*_*K*_ = 11. *±* 0 0.8 s^−1^ and a minimal contribution at 0.01 s appreciable with logarithmic binning (inset). These results suggest the generality of the frequency-dependent signal detection by talin folding dynamics, where the mechanical stability of the protein domain selects the responsive frequency regime. Additionally, we measured the folding probability of R3 WT in the presence and absence of external mechanical noise, observing no statistically significant difference (SI Appendix, Fig. S3). These data suggest that, similar to the mutant R3 IVVI, R3 WT filters out random mechanical fluctuations but responds to coherent force oscillations.

**FIG. 4.**
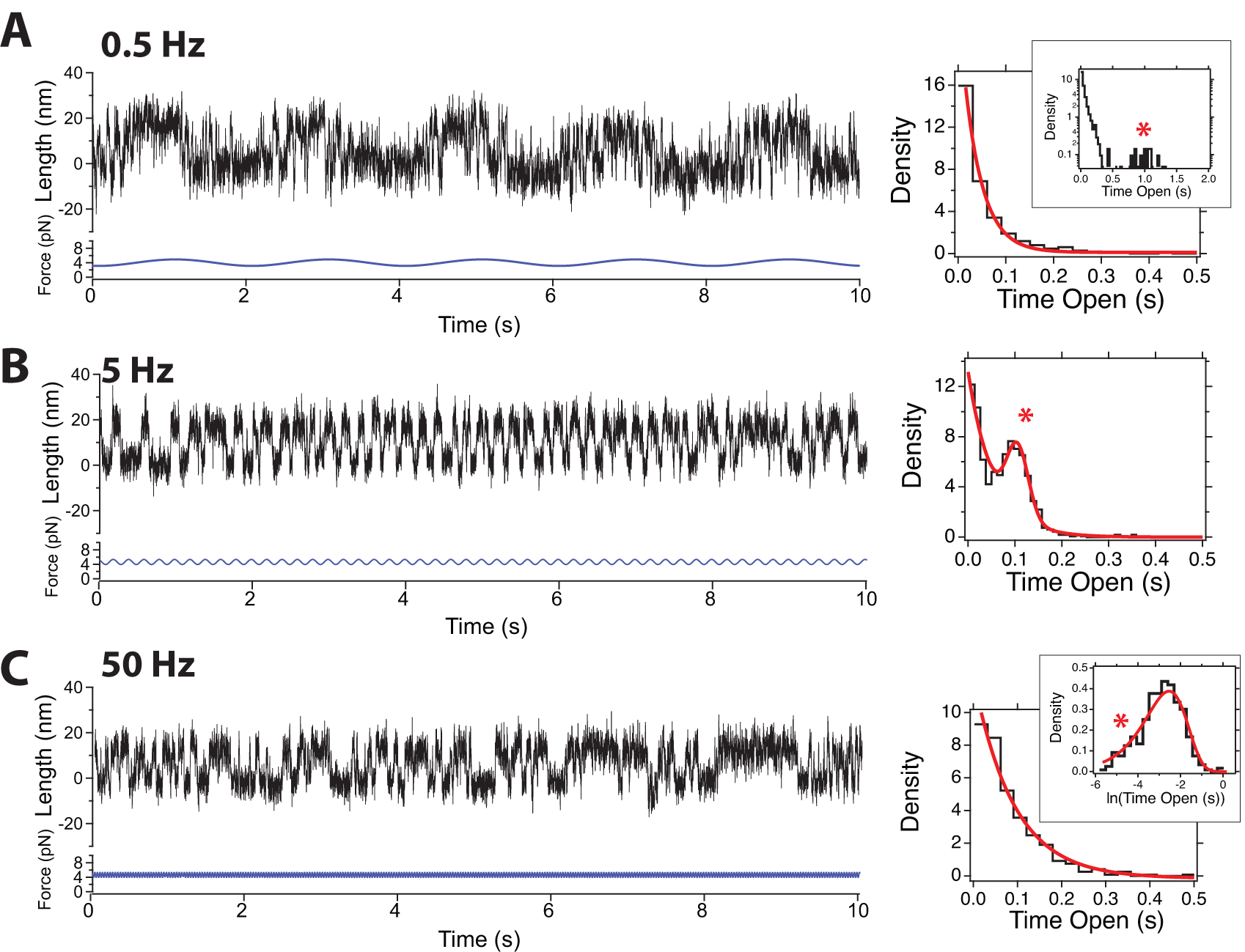
Response of the R3 WT domain under signals of different frequencies: (A, B, C) (left) 10 seconds-long fragments of magnetic tweezers recordings showing R3 WT under a sinusoidal force signal with an amplitude of 0.7 pN, and frequencies of 0.5, 5, and 50 Hz, respectively. (Right) Dwell-time histograms on the unfolded state, for each of these signals. The R3 WT domain has faster rates than R3 IVVI, and responds to higher frequencies. Histograms built with 715 (A), 2736 (B), and 1945 (C) transitions. Raw traces are shown, acquired at rates of 1-1.5 kHz.

### Stochastic Resonance in Talin Folding Dynamics

Our observations show a phase-locked response of talin folding dynamics as a mechanism of mechanical signal detection. This behavior resembles the physical phenomenon of stochastic resonance, by which non-linear systems exhibit an amplified response upon a weak input signal due to the presence of stochastic noise [28, 29]. Stochastic resonance has been demonstrated in very diverse contexts, such as climate dynamics [30], optical systems [31], DNA hairpin dynamics [32], or quantum systems [33] since it only requires three fundamental ingredients: a bistable system, a weak periodic input, and an intrinsic source of noise. All these three conditions are fulfilled by our system; the free energy landscape of the R3 domain is well-approximated by a symmetric double-well potential (SI Appendix, Fig. S4), and the thermal bath provides the intrinsic noise. Importantly, this intrinsic thermal noise is different from the external mechanical noise we used in Fig. 2, which was applied as an external random force on talin.

Signal detection by stochastic resonance typically refers to an increase in the signal-to-noise ratio (SNR) of the system output, which is the metric usually employed to characterize this physical phenomenon [28]. Indeed, the power spectrum of talin folding response under a periodic force perturbation (calculated on the idealized trace to remove the elastic contribution) exhibits a resonant peak at the driving frequency *f*_Ω_ (SI Appendix, Fig. S5). However, we can also characterize stochastic resonance by the shape of the dwell-time distributions, which, as discussed before, include a combination of stochastic and resonant transitions. Hence, these distributions can be modeled as the combination of an exponential distribution (stochastic), and a series of Gaussian peaks (resonant):

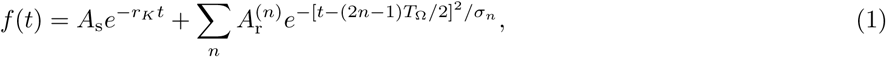

where *r*_*K*_ is the stochastic folding rate of talin (exponential contribution), *T*_Ω_ = 1*/f*_Ω_ the period of the driving force, and *A*_s_ and 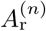 the amplitudes of the stochastic and resonant contributions, respectively. When the frequency is in the responsive regime, we observe several peaks at odd multiples of half the driving frequency ((2*n*− 1)*T*_Ω_*/*2, Fig. 3B). These contributions correspond to resonant transitions where one or more periods are missed but the transition occurs coherent with the driving signal at a future cycle. Hence, we can quantify statistically the fraction of resonant transitions *F*_R_ as the relative weight between the Gaussian distributions and the underlying exponential one. If the distribution is normalized (∫ *f* (*t*)*dt* = 1), *F*_R_ = 1 − *A*_s_*/r*_*K*_.

To demonstrate the response of talin over a broad range of frequencies, we carry out experiments on both the R3 IVVI and R3 WT domains under signals with *A* = 0.7 pN and frequencies ranging from 0.05 to 100 Hz. Figure 5A shows *F*_R_ as a function of the driving frequency *f*_Ω_. At frequencies above *r*_K_, the resonant peak can be buried in the shape of the exponential distribution, and it is not possible to directly determine its contribution. In such cases, we calculate the dwell-time histogram with logarithmic binning to separate fast timescales and estimate the relative weight from a logarithmic transform of Eq. 1 (Figs. S6 and S7). Similarly, we use the SNR to characterize stochastic resonance, measured from the power spectrum of the folding/unfolding time series (SI Appendix, Fig. S5). For both R3 IVVI and R3 WT, we observe the same dependence of the SNR on the driving frequency than that obtained using the fraction of resonant transitions, which illustrates the equivalence of both quantities to characterize stochastic resonance (SI Appendix, Fig. S5).

**FIG. 5.**
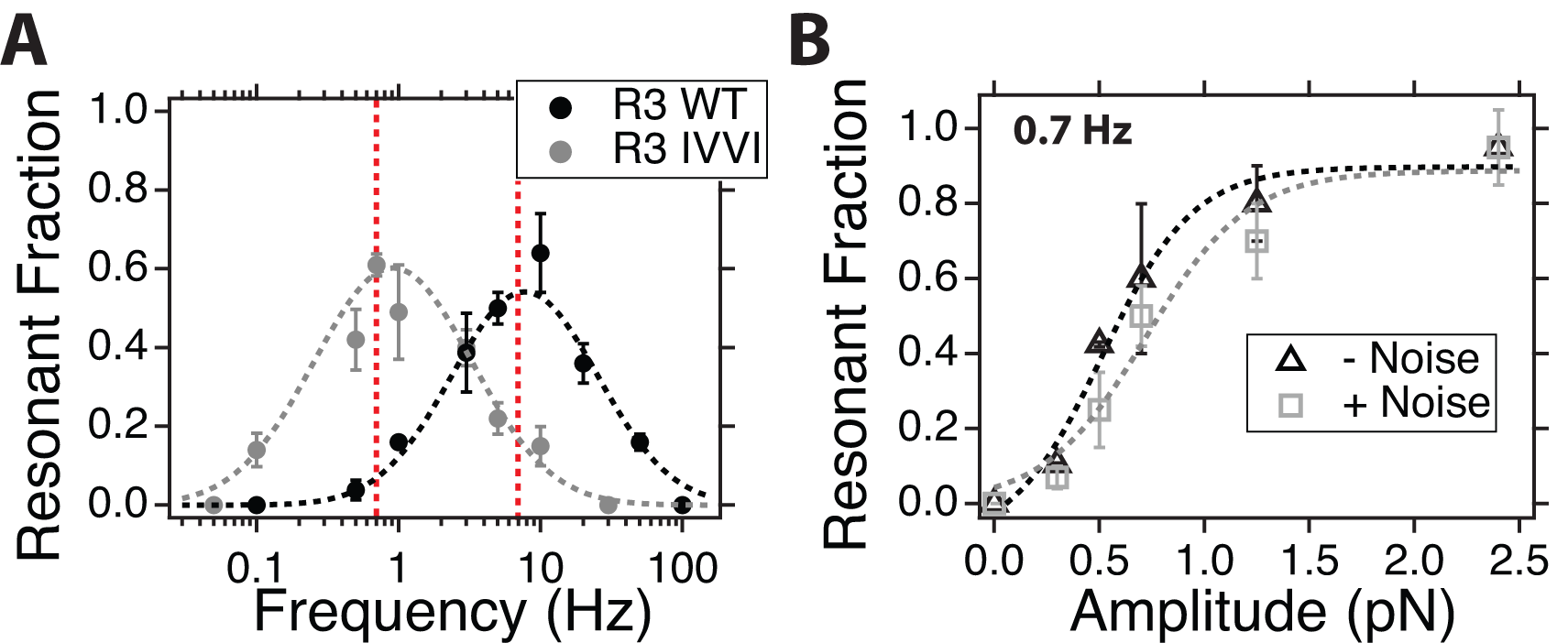
Quantitative characterization of the stochastic resonant dynamics by talin: (A) Fraction of resonant transitions for the R3 WT (black) and R3 IVVI (grey) domains. We characterize the level of resonant dynamics as the relative weight of the Gaussian-distributed transitions as measured from fits to the dwell-time distributions. Both domains show an optimal frequency range for signal detection around 10 Hz for R3 WT and 1 Hz for R3 IVVI. Higher frequencies are filtered, while lower ones show a majority of stochastic transitions due to the slow oscillation. Red lines show the theoretical resonance frequency, based on the natural folding rates at the coexistence force. The histograms and fits used to determine *F*_R_ are shown in SI Appendix, Figs. S6 and S7. (B) Fraction of resonant transitions as a function of the signal amplitude for a purely sinusoidal perturbation (black) and a sinusoid plus external mechanical noise (SD=2.8 pN; grey), measured for R3 IVVI at f=0.7 Hz. Dotted lines are sigmoidal fits to the data. Within statistical significance, there is no difference in talin response under either perturbation, which indicates that stochastic resonance is not affected by the addition of external mechanical noise to the coherent signal.

Figure 5A shows that talin operates as a mechanical bandpass filter, responding only over a narrow frequency range defined by the mechanical stability of the protein domain. For a bistable system, the optimal resonance frequency can be estimated as *r*_*K*_*/*2 [28] (red dotted line), which agrees with the experimental data. This resonance frequency is associated with the height of the intrinsic free energy barrier that separates the folded and unfolded states. Oscillating forces tilt the protein landscape towards the folded and unfolded states cyclically, increasing and decreasing the barrier periodically. If this periodic forcing occurs on a timescale compatible with the thermal-activated transitions, the state switching synchronizes with the perturbation. If the driving frequency is much faster than the intrinsic rates, the system responds only to the average barrier, filtering the oscillation; if the oscillation is too slow, the transitions are much more frequent than the timescale imposed by the external forcing.

Molecular kinetic processes measured with single-molecule techniques can be altered by the tethering system, which occasionally leads to artifactual effects arising from the interplay between the molecule and the dynamics of the pulling apparatus [34]. Hence, in order to test if our observations are an intrinsic property of the talin protein or are influenced by the magnetic probe, we conduct control experiments on R3 IVVI using magnetic beads of a larger size (M-450 with 4.5 *µ*m diameter, compared to the standard M-270 with 2.8 *µ*m diameter). These beads require a different chemical strategy to anchor our molecular constructs (SI Appendix, Supplementary Methods and Fig. S8). The experiments with the larger M-450 beads show no appreciable difference to those measured with the smaller M-270, despite its significant difference in size; R3 IVVI filters external mechanical noise of large bandwidth and gets optimally entrained under signals with a frequency of ∼1 Hz (SI Appendix, Figs. S9, and S10, S11). Hence, these data suggest that our observations are not altered by the tethering used in our experiments, likely because the diffusion coefficient of the magnetic beads is much higher than that of the elastic and folding response of talin, which, hence, become rate-limiting.

All the experiments in Fig. 5A were done using signals of constant amplitude, *A* = 0.7 pN, which is a moderate perturbation compared to the folding probability. In order to test the limit of signal detection, we measure the response of R3 IVVI at the resonance frequency *f*_Ω_ = 0.7 Hz and under different amplitudes (Fig. 5B). When the force signal spans the whole folding probability range (A>1 pN), the resonant response is saturated, and nearly all transitions are entrained with the periodic forcing. Under signals as small as 0.3 pN, we still observe an effect of resonant transitions of nearly 15%. This amplitude-dependent response is unaffected when the signal is buried in external mechanical noise (grey squares), which indicates that these random mechanical fluctuations do not influence the resonant properties exhibited by talin.This behavior is also reproduced with the R3 WT domain (Fig. S12), which suggests that both protein domains respond similarly to force signals that combine coherent oscillations and randomly fluctuating components. Although the amplitude of the periodic forcing enhances the level of signal detection, stochastic resonance does not arise simply from the magnitude of the oscillating forcing applied to the system. At an amplitude of *A* = 1.4 pN, where the response at the resonance frequency is saturated, we obtain the same frequency-response dependence as that at half that amplitude, indicating that the high selectivity of talin towards the driving force is amplitude-independent (SI Appendix, Fig. S13).

## DISCUSSION

Our data demonstrate that talin detects force oscillations by undergoing stochastic resonance. While stochastic resonance has been observed in biomolecules with very simple structures, such as RNA or DNA hairpins [32, 35], it is intriguing if this phenomenon is of generality among any protein under mechanical force, or if the talin R3 domain has some particularities due to its force-sensing function. Interestingly, the free energy landscape of both R3 IVVI and R3 WT at F_1*/*2_ has a symmetric bistable shape (SI Appendix, Fig. S4), which make them canonical models for stochastic resonance to occur [28]. By contrast, other proteins that have been characterized under force, such as protein L or the titin I91 domain, have very shallow force-dependencies of their unfolding rates, which leads to short distances to transition state from the folded state and very asymmetric folding landscapes [36, 37]. Hence, it is unclear if proteins with these characteristics would exhibit entrained folding dynamics, given that their folding landscapes do not fulfill the ideal conditions to undergo stochastic resonance.

Our most surprising result is the absence of a measurable effect of broadband external mechanical noise on the folding dynamics of talin. This is astonishing given that the mechanical noise amplitude of nearly 20 pN peak-to-peak used here dwarfs the 2 pN folding range of talin vastly. The physical basis of the external mechanical noise rejection mechanism is intriguing and contrasts sharply with the exquisite sensitivity shown by talin folding to periodic signals as low as 0.6 pN peak-to-peak. A possible interpretation of the effect of the external mechanical noise would be an effective increase of the temperature along the protein pulling axis; however, protein folding is very sensitive to temperature, and we do not observe any measurable effect of talin folding dynamics [38]. Although there is clearly much about the physics of proteins folding under force that remains poorly understood, our experiments are strongly suggestive that stochastic resonance could play an essential role in the biological function of talin as a mechanosensor.

Although initially formulated in the context of nonlinear physics, stochastic resonance has been demonstrated in a broad range of biological systems, with particular emphasis as a sensory mechanism in mechanoreceptors, like the crayfish hair cells [39], the cricket cercal system [40], or the vestibular and auditory system [6, 41]. Interestingly, in all these examples, signal transduction involves the activation of gated ion channels, which convert mechanical perturbations into electrophysiological signals. However, mechanotransduction also involves biochemical signaling, where force stimuli trigger downstream signaling pathways through a complex network of interacting proteins [3–5]. In this sense, it remains yet to be explored if stochastic resonance could also play a role in mechanotransduction pathways that involve ligand binding to force-bearing proteins instead of gating of mechanosensitive channels.

Mechanical signal transduction relies on the robust and finely-tuned response of molecular force sensors. Mechanical information is both encoded in the amplitude of the signal and its time-dependent evolution. Hence, both components must be accurately deciphered and interpreted by cellular force sensors. The auditory system is the paradigm of a mechanotransduction organ [7, 42, 43]. The hair cells are responsible for sound detection and transduction into electrical signals. Sound waves travel through the cochlea and induce vibrations on the basilar and tectorial membranes, which have a gradient of thickness and stiffness between the base and the apex. Hence, each frequency resonates at different positions along the membranes, which effectively decomposes the sound wave into its basic components that excite hair cells at different locations along the cochlea. The deflection of the hair bundles triggers the fast opening and closing of gated channels, ultimately converting the complex sound wave into nerve impulses that are interpreted by the brain. This process is highly nonlinear and has an excellent performance in the presence of noise, which indeed can induce deflections of the stereocilia over a hundred times larger than those caused by threshold stimuli [29]. Hence, stochastic resonance has often been discussed to operate as an adaptive mechanism for enhanced sensitivity in the naturally noisy hearing process [6, 41, 44, 45].

In a similar sense, cells constantly receive and transmit complex force stimuli between other cells and their environment, which are converted into biochemical signals that ultimately modulate gene expression. For example, durotaxis is directed by maintained fluctuating forces oscillating at frequencies 0.1 − 0.5 Hz that cell junctions apply to the extracellular matrix to sample its stiffness [15]. Additionally, protrusive podosomes generate oscillating forces that underpin their mechanosensing activity [46, 47]. At the protein level, single-cell measurements have demonstrated that talin experiences cyclic fluctuations with an amplitude that necessary involves folding and unfolding transitions [48]. All this recent evidence suggests the physiological relevance of force oscillations in mechanosensing; however, how cellular force sensors decompose these complex signals in a finely-tuned way remains mostly unknown.

Our results show that individual talin R3 domains detect force oscillations in a finely-tuned way by shifting its natural stochastic dynamics to a phase-locked folding response over a frequency range that is defined by the mechanical stability of the protein domain. The 13 talin-rod domains have a graded mechanical stability and respond to the amplitude of the force signal by unfolding hierarchically upon increasing loads [24, 25]. In the context of our findings, this could suggest that each talin bundle responds to complex force signals by undergoing stochastic resonance at different frequencies correlated with their unfolding force; weak domains would respond to fast frequencies while more stable ones would detect slow oscillations. Altogether, the talin rod would resonate at different segments over its extension, providing a mechanism for signal decomposition, analogous to that in the cochlear membranes.

Ultimately, the force on talin is transduced into a biochemical signal by the recruitment of binding proteins, which is regulated by the folding status of talin domains. There are at least 11 vinculin-binding sites buried in the hydrophobic core of the 13 talin bundle domains [16, 19]. Hence, vinculin binding is force-dependent, and maximal vinculin recruitment requires sufficient load on talin to unfold all the rod domains. By contrast, other ligands such as RIAM or DLC1 bind to folded talin bundles, and are inhibited by the amplitude of the force bore by talin, often competing antagonistically with vinculin for the same substrate [16, 49]. Therefore, the transduction of force stimuli into biochemical signaling by talin is controlled by force-dependent binding constants. Our data demonstrate that the emergence of resonant dynamics shifts the distribution of folding/unfolding transitions from stochastic to deterministic dynamics, which are characterized by regularly-spaced transitions. We propose that the change from random to deterministic folding kinetics in entrained talin domains plays a vital role in the binding kinetics of its ligands. If this proves to be correct, entrained folding dynamics may provide for a mechanism to convert complex mechanical signals into Fourier-decomposed biochemical signals along the talin mechanosensor.

## METHODS

subsection*Single-Molecule Experiments All our experiments were carried out in our custom-made magnetic tape head tweezers setup, detailed in [27]. The fluid chambers were built and functionalized, as described before [27, 50]. Experiments are carried out in HEPES buffer with 10 mM ascorbic acid (pH 7.3). The data are acquired at rates between 1 to 1.6 kHz with our custom-made software, written onto a binary file, and visualized in real-time with Igor Pro 8 (Wavemetrics). Single-molecule recordings are smoothed with a Savitzki-Golay algorithm with a 101-box size (R3 IVVI) or a 31-box size (R3 WT). Recordings are idealized for analysis with a double-threshold algorithm to detect the occupations on the folded and unfolded states automatically. Further details on experimental analysis are described in SI Appendix.

### Protein Expression and Purification

Polyprotein constructs are engineered using BamHI, BglII, and KpnI restriction sites in pFN18a restriction vector, as described previously [26]. Our protein construct contains the R3 IVVI, or R3 WT mouse talin domain, followed by eight titin I91 domains, and flanked by an N-terminal HaloTag enzyme and a C-terminal AviTag for biotinylation.

### Data Availability

All necessary data are available in the manuscript and SI Appendix.

