## Supplementary material for "Talin Folding as the Tuning Fork of Cellular Mechanotransduction": SI Appendix

### SUPPLEMENTARY METHODS

#### Magnetic Tweezers Setup

All the experiments are done on our custom-made magnetic tape head tweezers setup, as described in [1]. The instrument is built on top of an inverted microscope, using a 63x or 100x oil-immersion objective mounted on a piezo actuator (P-725; Physik Instrumente). The fluid chambers are illuminated using a collimated cold white LED (ThorLabs), and images are acquired using a CMOS Ximea MQ013MG-ON camera. Force is applied using a magnetic tape head (BRUSH 902836), positioned at 300 $\mu$m above the bottom glass of the fluid chamber, and controlled by the electric current applied to the tape head, calibrated as shown in [1] and maintained under feedback using a current-clamp PID controller. Individual molecules are tethered to superparamagnetic Dynabeads$^{TM}$ M-270 streptavidin beads, which allows to apply forces between 0 and 40 pN when using currents between 0 and 1000 mA. The data acquisition and control of the piezoelectric actuator is done using a multi-function DAQ card (NI USB-6361, National Instruments).

#### Fluid Chamber Preparation

All experiments are conducted in custom-made fluid chambers built by two sandwiched glass cover slides (Ted Pella) separated by a laser-cut parafilm pattern. The fluid chambers are functionalized with the HaloTag O4 ligand and the reference beads as described before [2]. Prior to the experiment, the fluid chambers are passivized for at least 3 hours using TRIS blocking buffer (20 mM Tris-HCL pH 7.4, 150 mM NaCl, 2mM MgCl2, and 1% w/v/ sulfhydryl blocked-BSA).

#### Protein Expression and Purification

The talin constructs are engineered using a combination of BamHI, BglII, and KpnI restriction sites in pFN18a restriction vector, as described previously [2]. Our protein construct for the magnetic tweezers experiment contains the R3 IVVI or R3 WT mouse talin domain, followed by eight I91 domains or two Spy-0128 domains as a molecular spacer (mechanically strong domains that do not unfold in the folding range of talin and, hence, do not interfere with talin dynamics), and flanked by an N-terminal HaloTag enzyme and a C-terminal AviTag for biotinylation. No appreciable difference between the I91 or the Spy0128 spacers has been observed. For purification purposes, a (His)6-tag is also present before the AviTag. BLR (DE3) or ERL competent cells are grown at 37°C, and the protein over-expression induced with 1 mM Isopropyl -D-1-thiogalactopyranoside (IPTG, Sigma) overnight at 25°C. Cells are re-suspended in 50 mM sodium phosphate buffer pH 7.0, 300 mM NaCl, 10% glycerol, and lysed. The proteins are purified from the lysate first with a Ni-NTA affinity resin, followed by size exclusion chromatography using a Superdex-200 HR column in 10 mM HEPES buffer pH 7.2, 150mM NaCl, 10% v/v glycerol and 1mM EDTA. The purified proteins are then concentrated to 50-100 $\mu$M and biotinylated in 50 mM Bicine buffer pH 8.3, 10 mM magnesium acetate, 10 mM ATP, 100 $\mu$M biotin and 2.5 $\mu$g biotin ligase BirA enzyme (Avidity), at 40°C for 4 hours or overnight at 4°C. For the M-450 control experiments (see below), the C-terminal of molecular construct contains a SpyTag peptide instead of the AviTag.

#### Single-Molecule Measurements

All experiments are conducted in HEPES buffer (Hepes 10 mM pH 7.2, NaCl 150 mM, EDTA 1 mM) containing 10 mM ascorbic acid (pH 7.4) to minimize oxidative damage. The protein is incubated on the fluid chamber for $\sim$30 minutes to a concentration of 1-5 nM, and then washed using the measuring buffer to remove the non-anchored molecules. The M-270 streptavidin beads are passivized in TRIS blocking buffers for at least 3 hours before the start of the experiment. A volume of $\sim$20 $\mu$L of magnetic beads is added to the chamber and left for $\sim$2 minutes at 0 pN to incubate. The molecule-searching process is done at a force of 4 pN for R3 IVVI and at a force of 3 pN for R3 WT, forces low enough to ensure that all proteins are folded, but applying some tension to prevent the settlement of the beads. Individual molecules are identified through a fingerprint pulse consisting on a reversible force ramp between the resting force and 20 pN with a duration of 10-20 seconds. Individually tethered molecules show a single unfolding step and a single refolding step of $\sim$ 17 nm at a force of $\sim$10 pN for R3 IVVI or $\sim$5 pN for R3 WT. Once the fingerprint is observed the desired measurements starts.

### Application of Mechanical Signals

The use of the magnetic tape head allows to subject individual molecules to force signals that change quickly in time. The magnetic tape head, controlled by a current-clamp PID circuit, has a bandwidth of ∼10 kHz, as characterized in [1]. For the application of mechanical signals, we use a high speed DAQ (NI USB 6289) whose output is added to that of the acquisition DAQ card. The high speed DAQ generates the time-changing signals built by an independent custom-written software, while the main DAQ card generates the offset forces (*e.g.* when applying a sinusoidal force signal of 1 Hz, an average of 9 pN and an amplitude of 0.7 pN, the high speed DAQ outputs a sinusoidal signal of 1 Hz and 0.7 pN of amplitude, with average of 0, which is summed to a constant force of 9 pN applied by the main DAQ). The high speed DAQ generates signals with a 60 seconds duration that are sampled at a 100 kHz rate and buffered to be looped for the desired duration in time. The noise signals are generated as pseudo-random numbers with a Gaussian distribution of the chosen standard deviation using the Box-Muller algorithm.

### Control Experiments with the M-450 beads

Dynabeads$^{TM}$ M-450 tosylactivated superparamagnetic beads are employed to test the influence of the magnetic probe on talin resonant dynamics. These beads have a larger radius than the M-270 (2.3 $\mu$m versus 1.4 $\mu$m) so their diffusion coefficient is approximately half that of the M-270. There are not commercially available M-450 beads coated with streptavidin, which requires using a different functionalization strategy for anchoring the individual talin constructs (Fig. S9) based on HaloTag chemistry and the SpyCatcher-SpyTag split-protein technique. We generate two different protein constructs: (HaloTag)-(R3IVVI)-(Spy0128)$_2$-SpyTag, and (SpyCatcher)-(HaloTag). The first construct is anchored to the glass surface using the usual HaloTag chemistry. The M-450 beads are functionalized with the HaloTag ligand with a similar protocol to that used for the surface. The molecular construct is closed by incubation of the beads in the fluid chamber for ∼ 5 minutes, which allows the formation of the intermolecular isopeptide bond between the glass-anchored SpyTag and the bead-anchored SpyCatcher. Hence, this strategy allows for a double-covalent anchoring of single protein constructs.

To observe the effect of force signals on R3 IVVI dynamics measured with the M-450 magnetic probes, we look for the electric current at which talin populates the folded and unfolded states equally ($F_{1/2}$), which corresponds to 135 mA (in contrast to 320 mA with the M-270). We use an amplitude of 10 mA for the sinusoidal signals, which corresponds to 0.7 pN, assuming that 135 mA is 9 pN ($F_{1/2}$) and a linear force-current relation in the low force range, as shown in [1].

### Single-Molecule Data Analysis

*a. Data storage and visualization:* Our data is stored in real-time as a four-column binary file containing the time mark (s), the protein extension (nm), the force (pN), and the electric current measured through the head (mA). We visualize these data in real time using a custom-written software in Igor Pro (Wavemetrics). Data is acquired at sampling rates between 1,000 and 1,600 Hz, depending on the size of the region of interest and smoothed with a 4$^{th}$-order Savitzky-Golay filter using a box size of N=101 for R3 IVVI or N=31 for (R3 WT), due to its faster dynamics. The folded/unfolded states of talin are automatically detected using a double threshold algorithm (Fig. S2).

*b. Fraction of resonant transitions:* The fraction of resonant transitions (Fig. 5, main text) we use to characterize stochastic resonance is obtained from fits to Eq. 1 (main text) as $F_R = 1 - A_s/r_K$ (see Figs. S7, S8, and S11). Fits are done forcing the Gaussian components to be centered at odd multiples of the half-period of the driving signal.

*c. Signal-to-noise ratio:* Alternatively, we use the signal-to-noise ratio (SNR) to characterize the resonant response of talin folding dynamics (Fig. S6). In order to remove the contribution of the polymer elasticity, we calculate the power spectrum on the idealized trace (binary open-closed time-series). The power spectrum $S(f)$ is defined as the scaled magnitude by calculating a fast Fourier Transform from a Discrete Fourier Transform of the idealized trace time-series by using a prime factor decomposition algorithm. The power spectrum consists of the combination of a background spectral density, and resonant spikes centered at odd multiples of the driving frequency, since the input signal is a binary time-series of open-closed states (Figure S6A). To calculate the SNR, we fit a Lorentzian at the vicinity of the resonant peak magnitude at the driving frequency, and calculate the ratio between the peak and the baseline (Figure S6B). The error bars are calculated from the uncertainty of the fitted parameters.

### SUPPLEMENTARY FIGURES

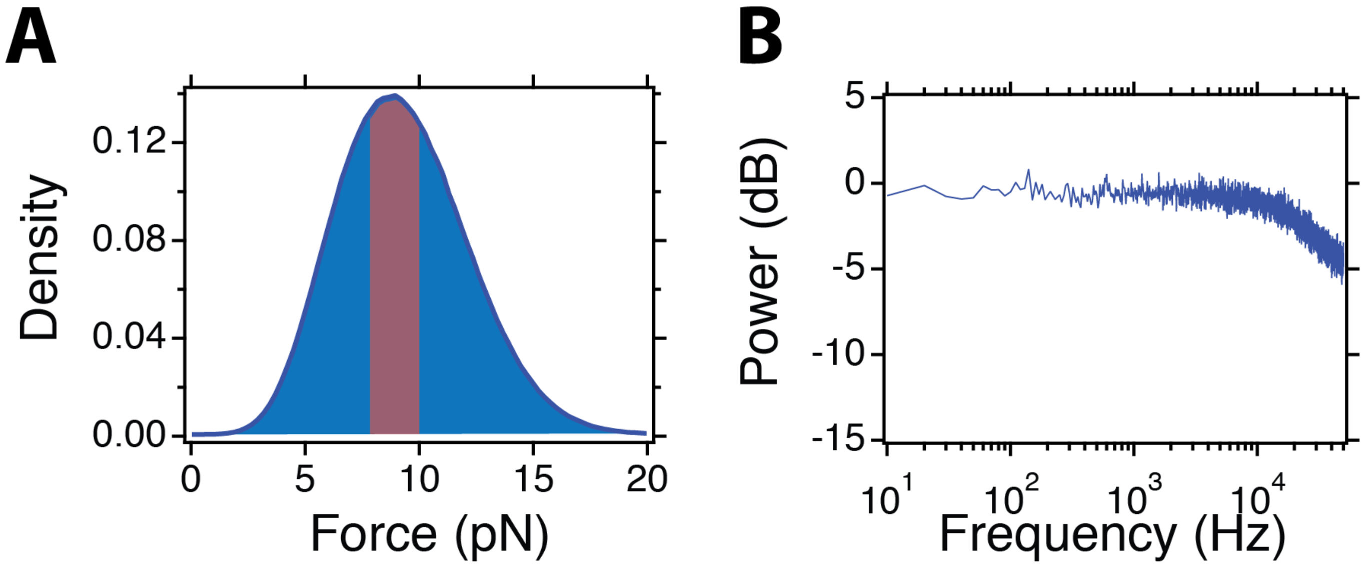


FIG. 1. **Distribution and power spectrum of the external mechanical noise**: **(A)** Force distribution of the external mechanical noise. The external mechanical noise is generated as described in the Supplementary Methods. To test the actual force signal we are applying, we measure the electric current across the magnetic tape head using a high-speed DAQ running at 100 kHz, and convert it to force as described in [1]. The external mechanical noise has a Gaussian distribution with an average of 9 pN and a standard deviation of 2.8 pN. The red shaded region is the width of the folding probability of R3 IVVI, which illustrates that the external mechanical noise fluctuates over a force range that greatly exceeds that of R3 IVVI folding. Equivalent noise properties are used on the R3 WT, but with an average of 5 pN. **(B)** Power spectral density of the external mechanical noise. The mechanical noise has a flat power spectrum up to the head bandwidth ($\sim$ 10 kHz).

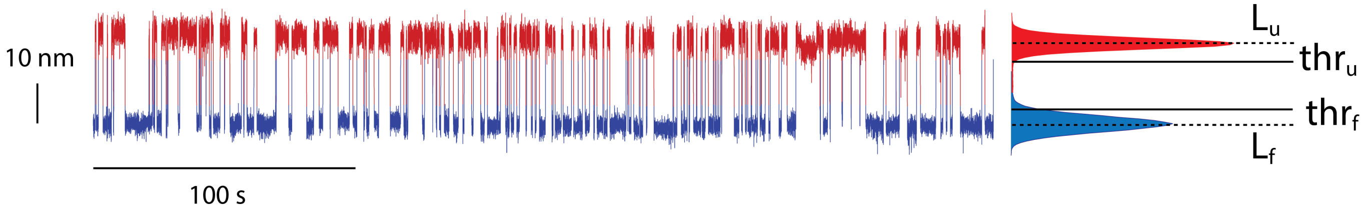


FIG. 2. **Illustration of the double-threshold algorithm for automatic detection of the talin folding status**: Typical R3 IVVI recording with the folded (blue) and unfolded (red) states identified using the double-threshold algorithm. From the histogram of the protein extension time series, we determine the two thresholds based on the position of the histogram peaks. The status of the protein is followed over time and changed if the opposite threshold is overcome. *i. e.* Assuming that the protein is folded, its status changes to unfolded if the threshold $thr_U$ is crossed, and vice-versa. This method is more robust than using a single threshold, and particularly useful when analyzing entrained recordings, where the elastic component of the protein changes following the signal. From the time series of the idealized trace (open-closed binary time series), we calculate all metrics used to characterize the folding properties of R3 IVVI and R3 WT upon force signals (*e. g.* dwell time distributions, power spectrum, ...).

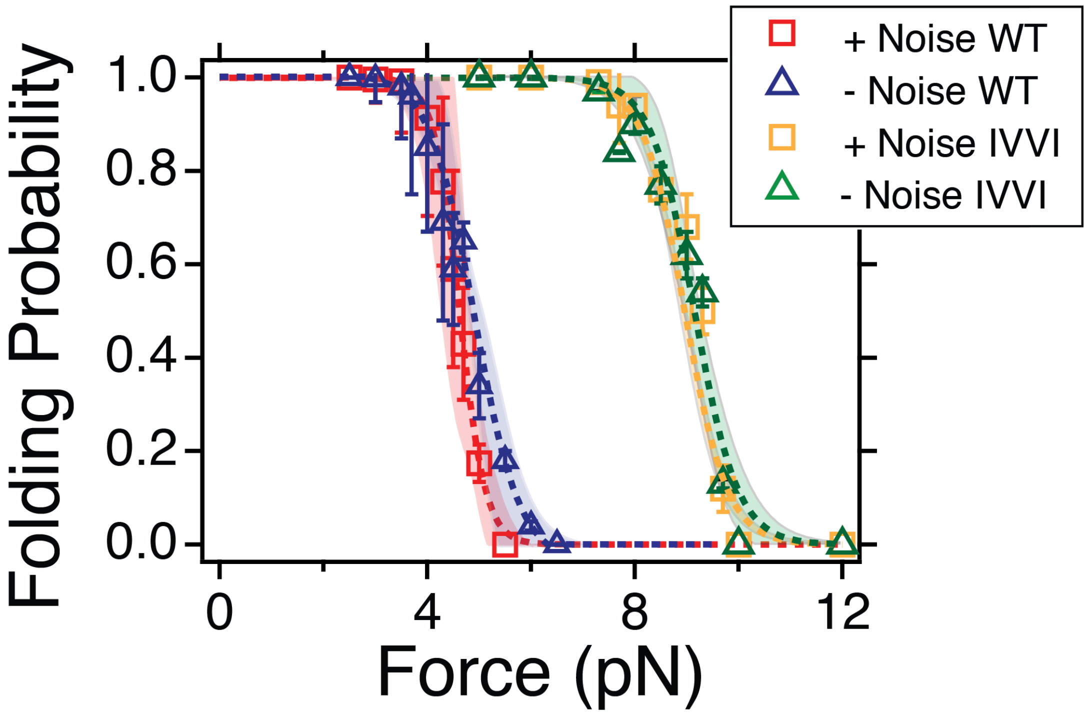


FIG. 3. **Folding probability of R3 WT (red and blue) and R3 IVVI (orange and green) in the presence (squares) and absence (triangles) of external mechanical noise**. Error bars are the SEM of the folding probabilities, calculated by propagating the SEM of the average residence times. Shaded regions show the 95% confidence contour, calculated from weighted sigmoidal fits using the Levenberg-Marquardt least orthogonal distance method. With a 95% confidence level, the folding probabilities overlap, suggesting that the mechanical noise produces no measurable effect on the R3 WT or R3 IVVI folding dynamics. The coexistence forces obtained from the fits overlap within error bars: R3 WT: $F_{1/2} = 4.91 \pm 0.07$ pN (- Noise); $F_{1/2} = 4.65 \pm 0.37$ pN (+Noise); R3 IVVI: $F_{1/2} = 9.15 \pm 0.15$ pN (- Noise); $F_{1/2} = 9.14 \pm 0.12$ pN (+Noise).

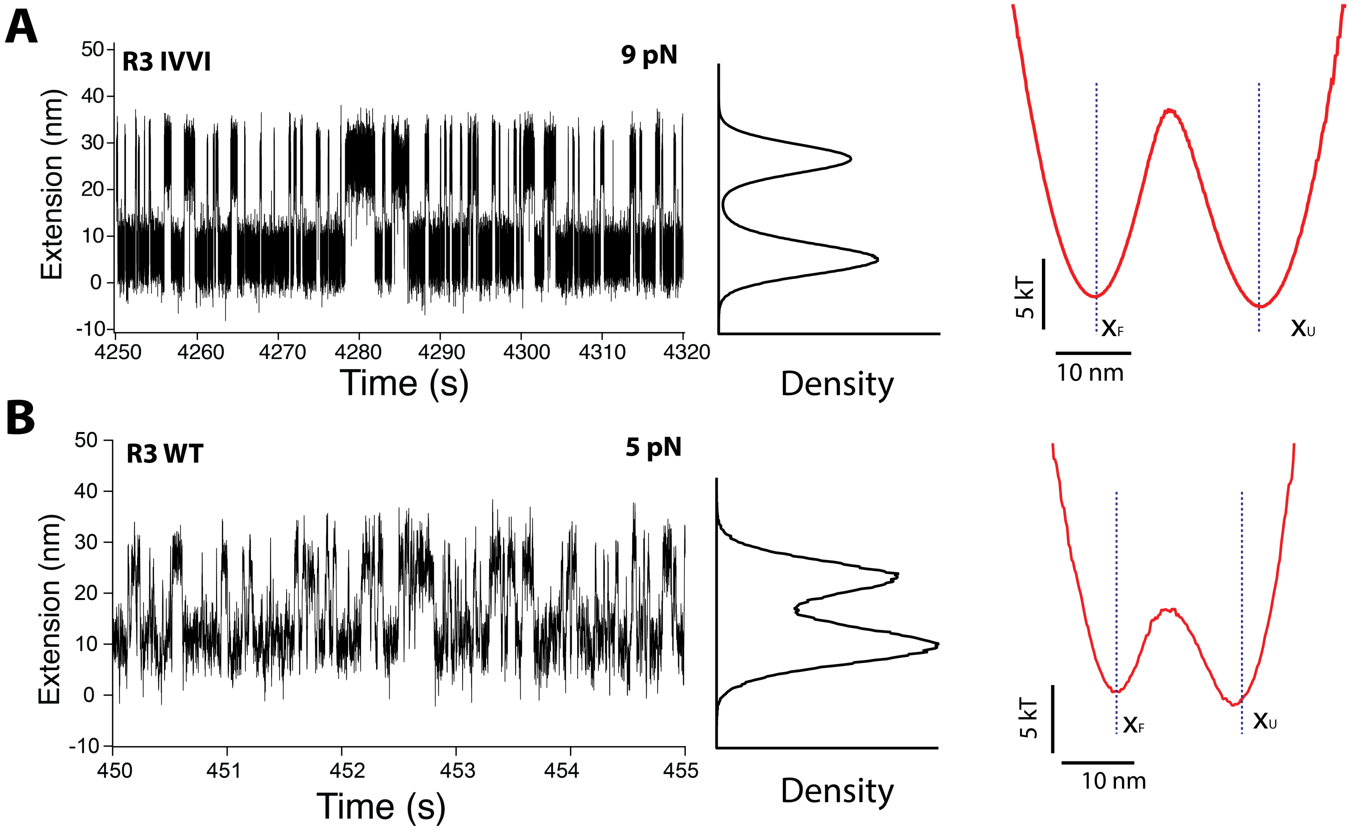


FIG. 4. **Free Energy Landscape of R3 IVVI (A) and R3 WT (B) at their respective $F_{1/2}$:** Fragments of trajectories for both R3 variants at their respective $F_{1/2}$, showing the reversible folding dynamics (left). We estimate their free energy projections along the pulling coordinate using the Boltzmann inversion formula $F(x) = -kT \ln P(x)$, being $x$ the pulling coordinate and $P(x)$ the distribution density of the protein dynamics along $x$, shown in the central panels. The free energy profiles of both proteins has the shape of a symmetric double-well potential, with very different free energy barrier heights that indicates their different kinetics (right). Hence, their folding dynamics under force can be well modeled as a bistable system, a condition required for stochastic resonance. Landscapes reconstructed from a single trajectory containing N=4506 transitions for R3 IVVI and N=2377 R3 WT.

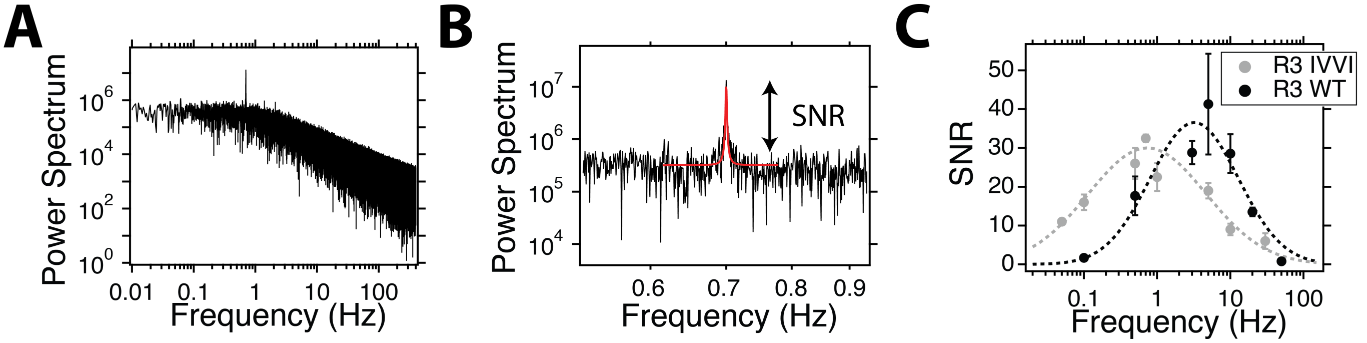


FIG. 5. **Signal-to-noise ratio quantification of stochastic resonance**: **(A)** Power spectrum of the R3 IVVI domain under a sinusoidal force signal of 0.7 Hz. The power spectrum is calculated on the time-series of the idealized trace (binary open-close time-series) to remove the contribution of the polymer elasticity and account only for the folding dynamics. **(B)** Detail of the peak region, highlighting the calculation of the signal-to-noise ratio. A Lorentzian is fitted in the vicinity of the peak. The SNR is calculated from the fit as the ratio between the peak's magnitude and the baseline. **(C)** SNR as a function of the frequency of the driving force signal for the R3 IVVI (grey) and R3 WT (black). A log-normal peak is fitted for visualization purposes (dotted lines). We obtain the same qualitative behavior as when using the resonant fraction; talin folding dynamics show a frequency-dependent resonant response, at frequencies of $\sim 1$ Hz for R3 IVVI and $\sim 10$ Hz for R3 WT.

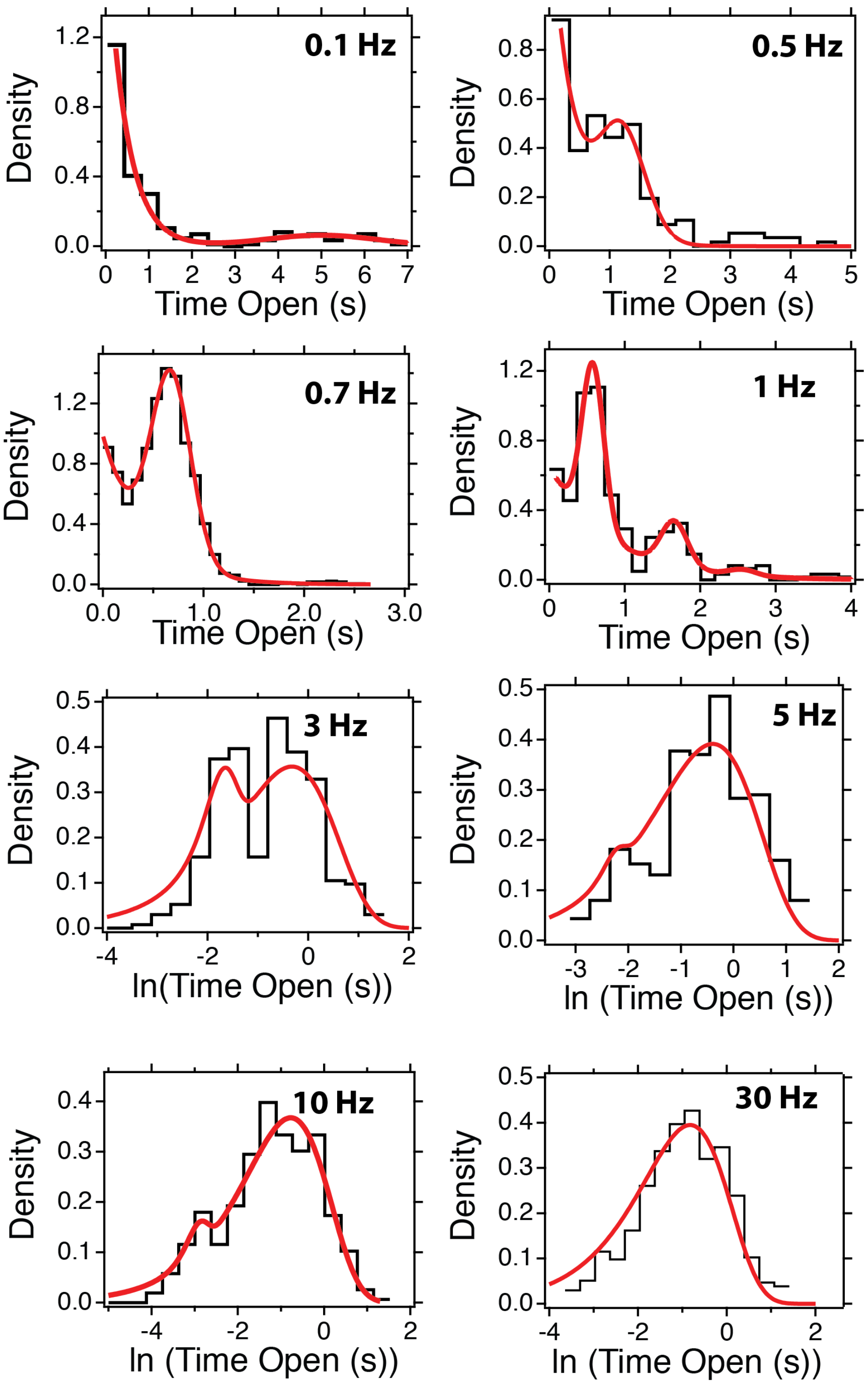


FIG. 6. **Dwell-time histograms of R3 IVVI obtained under sinusoidal signals of 0.7 pN of amplitude**: Red solid lines are fits to Eq. 1 in the main text. Above 3 Hz, the histograms are calculated with logarithmic binning and the fitting function transformed accordingly. Data obtained on a minimum of three different molecules per frequency and a number of transitions: f=0.1 Hz, N=223; f=0.5 Hz, N=192; f=0.7 Hz, N=1822; f=1 Hz, N=337; f=3 Hz, N=397; f=5 Hz, N=363, f=10 Hz, N=414; f=30 Hz, N=696.

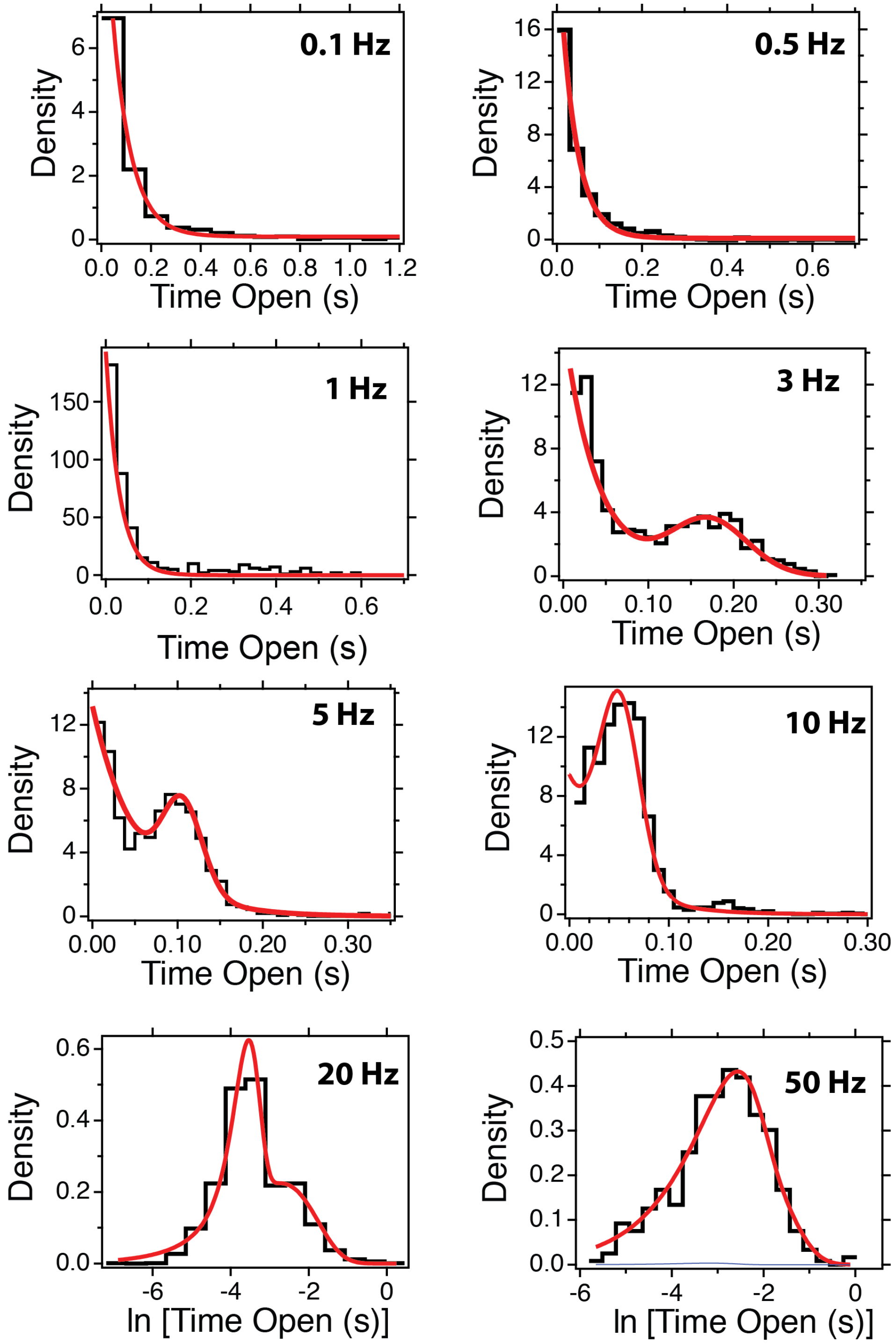


FIG. 7. **Dwell-time histograms of R3 WT obtained under sinusoidal signals of 0.7 pN of amplitude**: Red solid lines are fits to Eq. 1 in the main text. Above 20 Hz, the histograms are calculated with logarithmic binning and the fitting function transformed accordingly. Data obtained on a minimum of three different molecules per frequency and a number of transitions: f=0.1 Hz, N=543; f=0.5 Hz, N=705; f=1 Hz, N=940; f=3 Hz, N=1043; f=5 Hz, N=2736; f=10 Hz, N=1933, f=20 Hz, N= 1946; f=50 Hz, N=409.

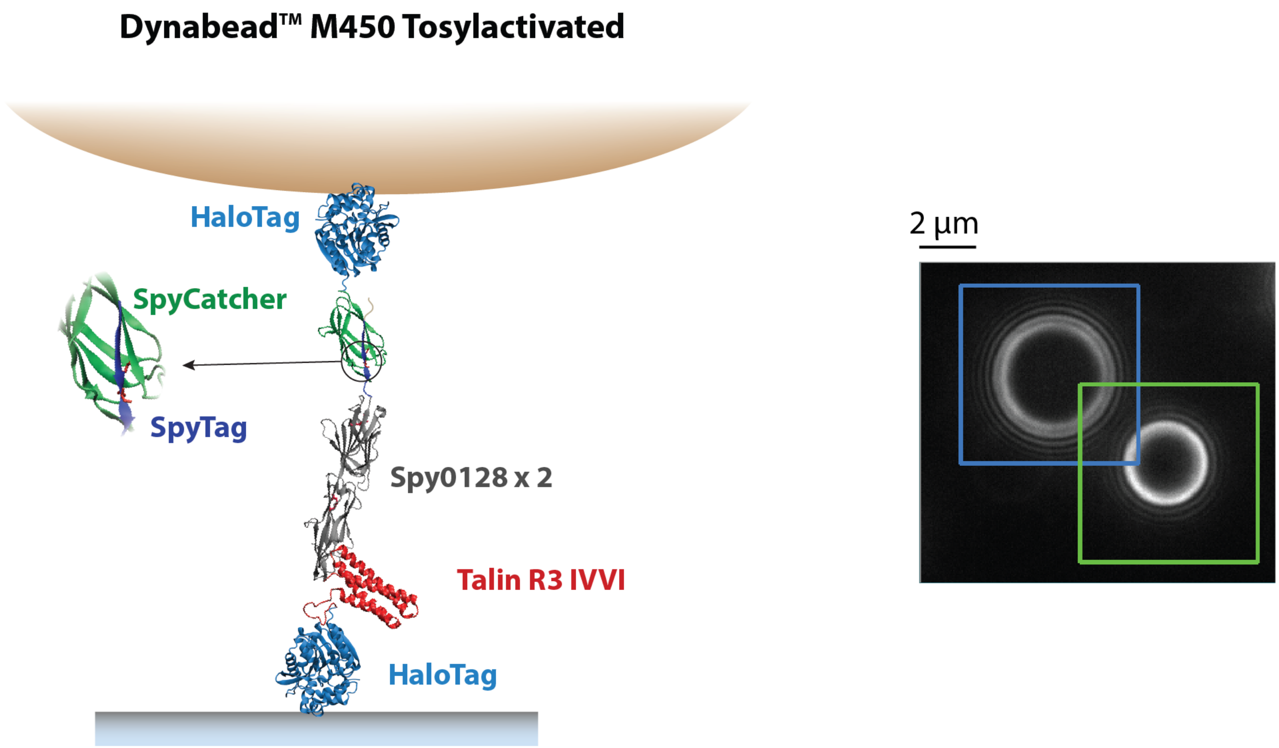


FIG. 8. **Single-molecule tethering to Dynabead$^{TM}$ M-450 tosylactivated superparamagnetic beads**: (Left) Schematics of the molecular construct employed to anchor single talin domains to the M-450 beads. These 4.5 $\mu$m diameter beads are not commercially available with a streptavidin coating; hence, we use a different chemical strategy to anchor single-molecule constructs using a double-covalent approach by combining HaloTag chemistry and split-protein techniques. The M-450 beads are incubated with HaloTag-SpyCatcher, while our protein construct containing the R3 IVVI domain is flanked by a HaloTag for covalent anchoring to the glass surface and a SpyTag peptide. After adding the M-450 beads to the fluid chamber, the protein construct and M-450 beads are covalently anchored through a isopeptide bond formed between the SpyTag and the SpyCatcher, which close the molecular tether. (Right) Photograph of an M-450 bead (blue square) close to a 3.5 $\mu$m diameter reference bead (green square.)

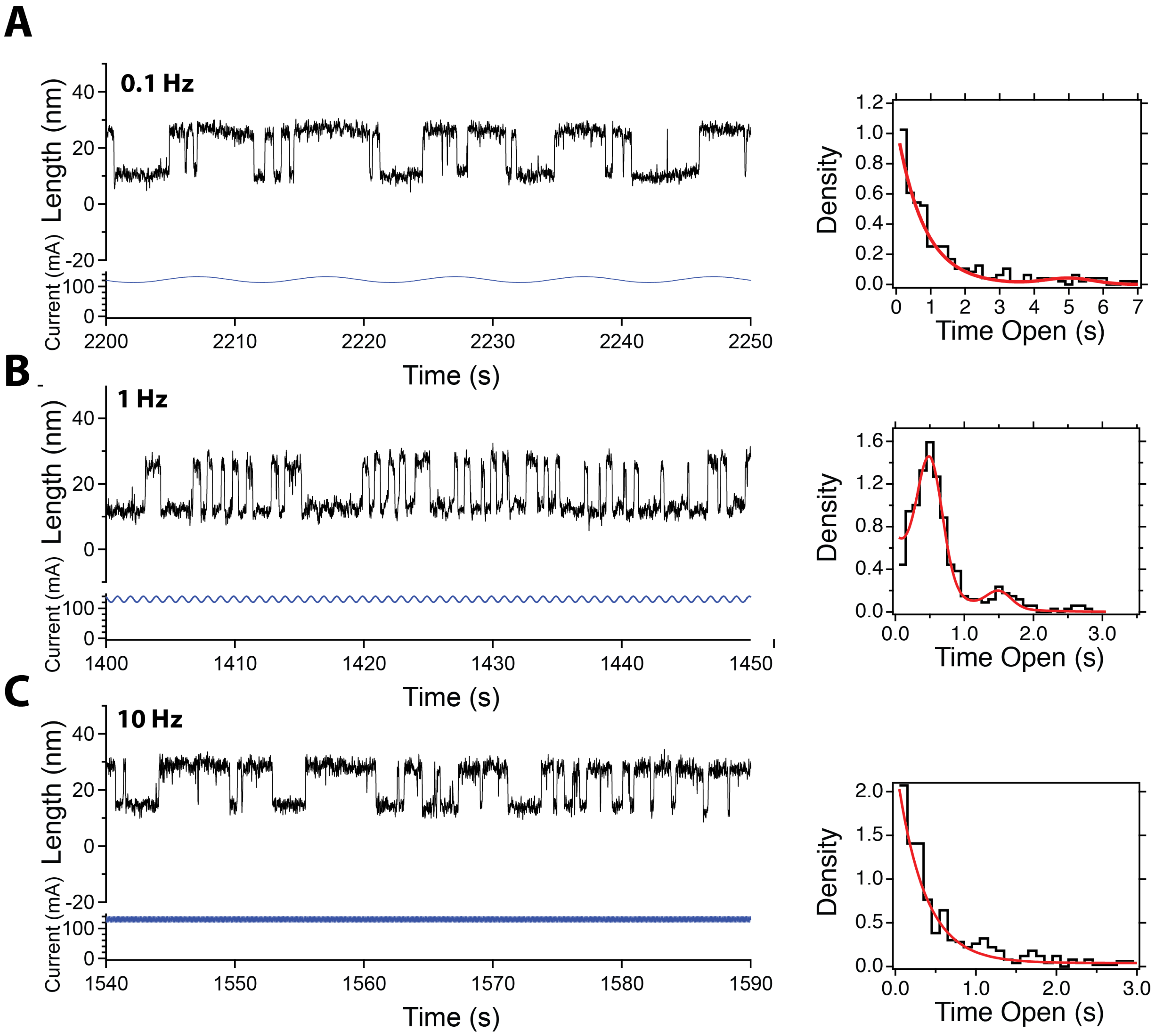


FIG. 9. **R3 IVVI dynamics under sinusoidal force signals measured with M-450 superparamagnetic beads**: Talin dynamics are optimally entrained at frequencies in the ∼1 Hz range (B), while we only observe a meager synchronized contribution at 0.1 Hz and 10 Hz. No appreciable differences are observed between in the experiments conducted with the M-270 or M-450 beads.

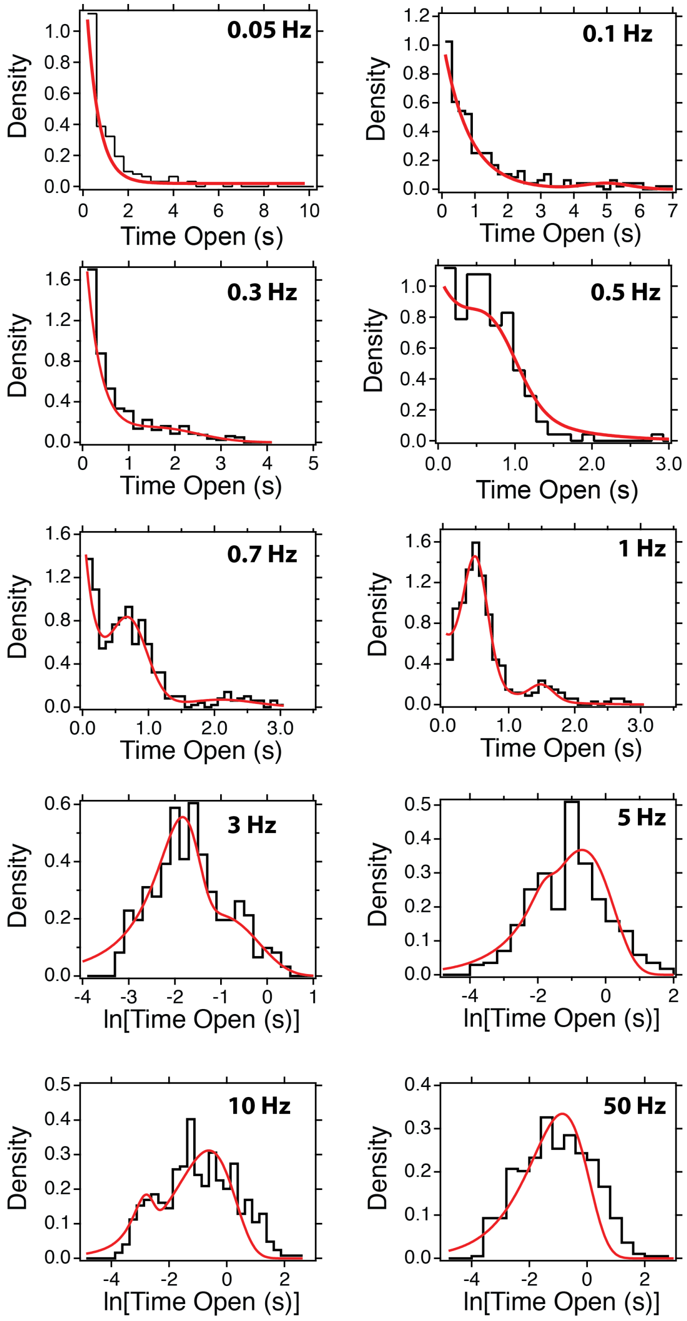


FIG. 10. **Dwell-time histograms for R3 IVVI under sinusoidal signals of 0.7 pN of amplitude measured with the M-450 beads**: Red lines are fits to Eq. 1 in the main text. Histograms built with three different individual molecules and the following number of transitions: f=0.05 Hz, N=155; f=0.1 Hz, N=239; f=0.3, N=405; f=0.5 Hz, N=161; f=0.7 Hz, N=496; f=1 Hz, N=339; f=3 Hz, N=306; f=5 Hz, N=477; f=10 Hz, N=497; f=50 Hz, N=483.

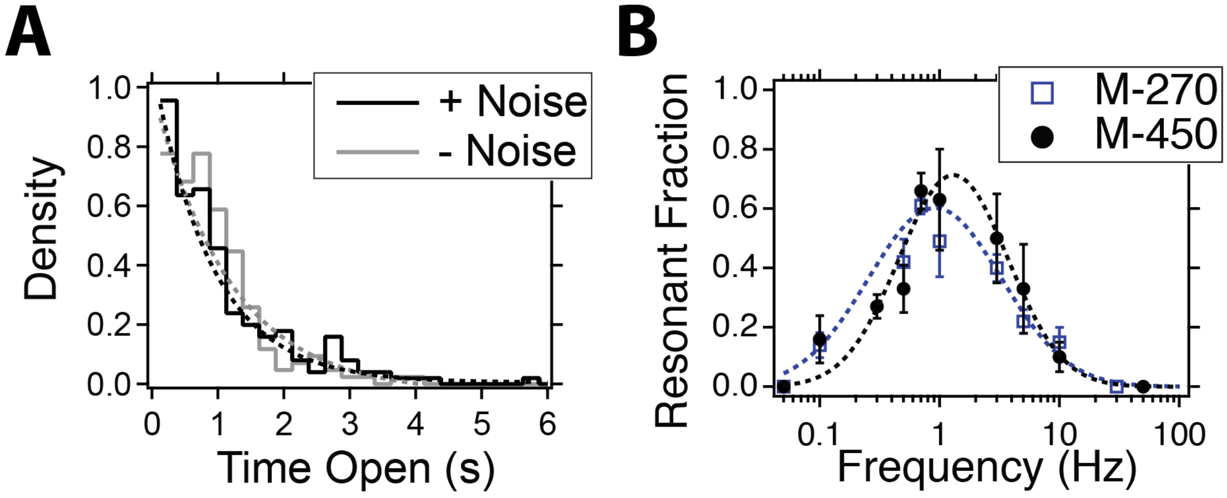


FIG. 11. **Talin response under force signals measured with the M-450 beads**: **(A)** Dwell-time histogram of R3 IVVI at $F_{1/2}$in the presence (black) and absence (grey) of external mechanical noise measured with the M-450 beads. The properties of the mechanical noise are equivalent to those shown in Fig. S1. As demonstrated with the M-270 beads, talin filters mechanical noise, which suggests that this property is independent of the employed tethering. The obtained average dwell times ($\tau$ = 0.90±0.07 s, + Noise; $\tau$ = 1.17±0.17 s. - Noise) are equivalent to those measured with the M-270. Histograms built with N=210 transitions (+ Noise) and N=170 transitions (- Noise). **(B)** Fraction of resonant frequencies for R3 IVVI under sinusoidal signals of 0.7 pN amplitude measured with the M-450 (blue) and M-270 (black) beads. The results obtained with the M-450 beads overlap within error bars with those measured with the M-270 beads, which suggests that stochastic resonance is an intrinsic property of talin, which is unaffected by the single-molecule tethering. Dotted lines are log-normal fits used for visualization reasons.

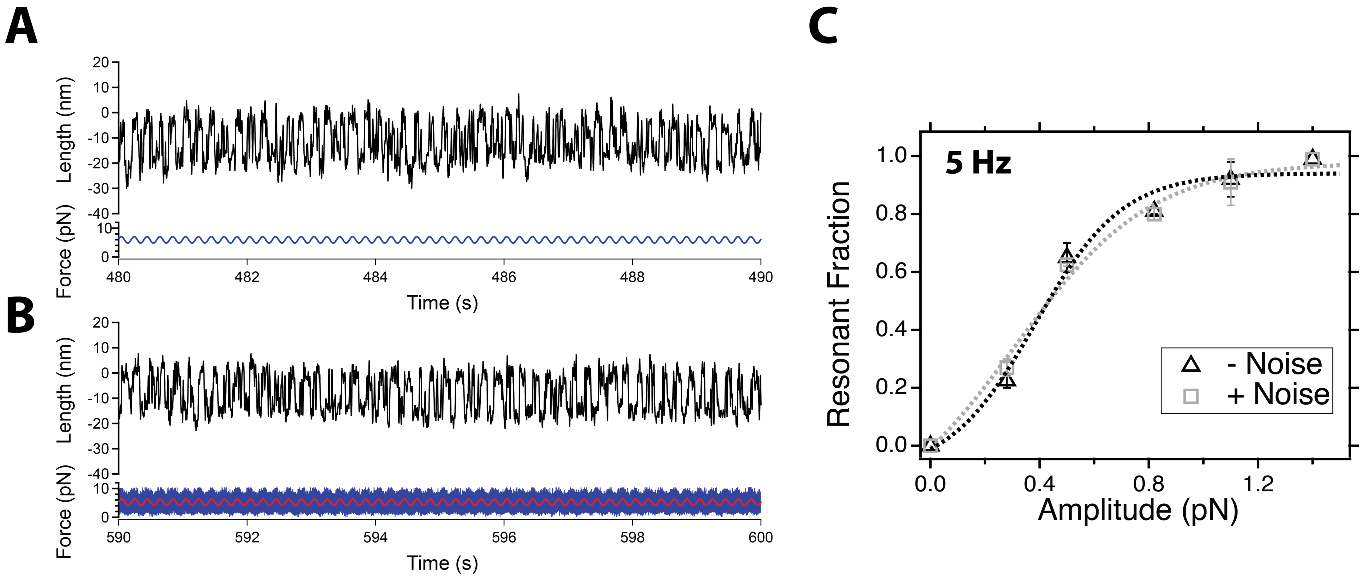


FIG. 12. **Compared folding dynamics of R3 WT under purely oscillatory and noisy oscillatory signals**: **(A)** Dynamics of the talin R3 WT domain under a sinusoidal signal of 5 Hz, 0.5 pN of amplitude, and an average of 5 pN. **(B)** Dynamics of the talin R3 WT domain under a force signal composed of the same sinusoidal used in (A), summed to external mechanical noise with a standard deviation of 2.8 pN. **(C)** Fraction of resonant transitions measured for R3 WT as a function of the amplitude of the oscillatory perturbation of 5 Hz, using a purely sinusoidal signal (black triangles) and a sinusoidal submerged in white mechanical noise with 2.8 pN standard deviation (grey squares). Dotted lines are sigmoidal fits to the data. There is no statistically significant difference in the response of R3 WT, which indicates that stochastic resonance is not affected by the addition of external mechanical noise to the coherent signal.

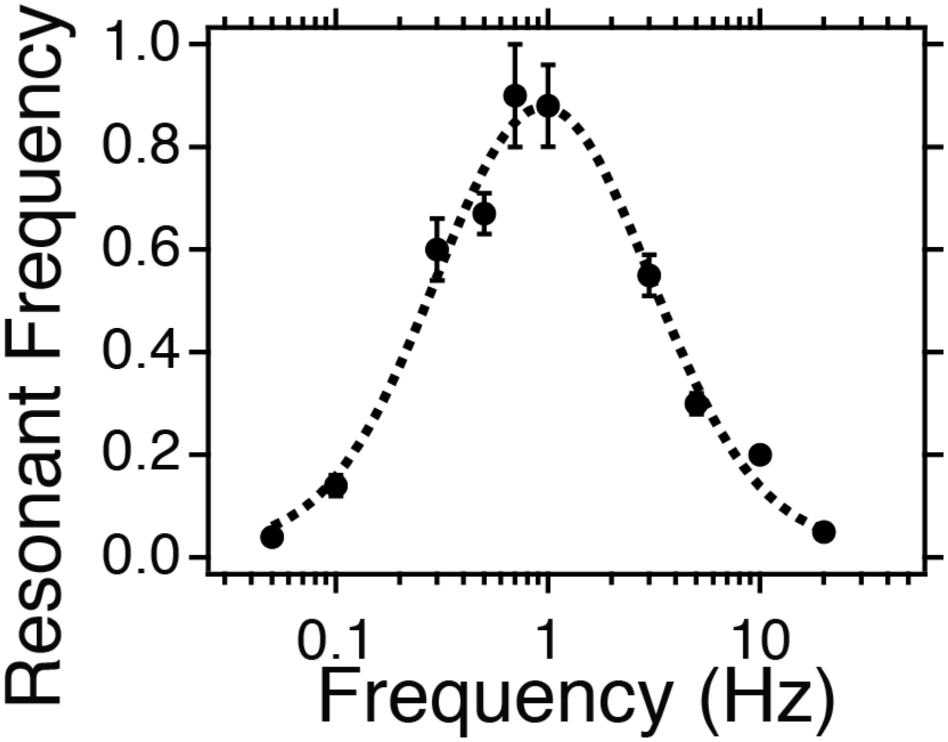


FIG. 13. **Fraction of resonant transitions under a sinusoidal signal of 1.4 pN of amplitude measured for R3 IVVI**: The larger amplitude increases the number of resonant transitions to nearly 100% in the ∼1 Hz range. However, we observe a strong frequency-dependent behavior, which indicates that the stochastic resonance phenomenon is not dependent on the signal amplitude, and that talin only responds on a narrow frequency range.
